# Assessing the predictive ability of computational epitope prediction methods on Fel d 1 and other allergens

**DOI:** 10.1101/2023.06.01.543222

**Authors:** Hyeji Kwon, Soobon Ko, Kyungsoo Ha, Jungjoon K. Lee, Yoonjoo Choi

## Abstract

While computational epitope prediction methods have found broad application, their use, specifically in allergy-related contexts, remains relatively less explored. This study benchmarks several publicly available epitope prediction tools, focusing on the allergenic IgE and T-cell epitopes of Fel d 1, an extensively studied allergen. Using a variety of tools accessible via the Immune Epitope Database (IEDB) and other resources, we evaluate their ability to identify the known linear IgE and T-cell epitopes of Fel d 1. Our results show a limited effectiveness for B-cell epitope prediction methods, with most performing only marginally better than random selection. We also explored the general predictive abilities on other allergens, and the results were largely random. When predicting T-cell epitopes, ProPred successfully identified all known Fel d 1 T-cell epitopes, whereas the IEDB approach missed two known epitopes and demonstrated a tendency to over-predict. However, when applied to a larger test set, both methods performed only slightly better than random selection. Our findings show the limitations of current computational epitope prediction methods in accurately identifying allergenic epitopes, emphasizing the need for methodological advancements in allergen research.

## Introduction

The cat allergy is one of the most common pet allergies, affecting approximately 20% of the global population[1]. The severity of cat allergy can vary, ranging from mild symptoms like coughing, pruritus, and skin eruption to potentially life-threatening reactions such as anaphylaxis. The primary allergen responsible for the cat allergy is Fel d 1, present in the fur and dander of domestic cats. It is estimated that approximately 90% of cat allergies are caused by Fel d 1[2].

Fel d 1 is primarily secreted by epithelial cells and the salivary glands of cats, and it remains on their haircoats during grooming. One notable characteristic of Fel d 1 is its high thermal stability, allowing it to persist on cat hair. It can associate with small airborne particles, spreading throughout the surroundings and leading to allergic reactions in susceptible individuals. In fact, a study reported that Fel d 1 was detected in 99.9% of households in the United States, highlighting its ubiquity and potential for exposure [2–3].

Although the function of Fel d 1 remains largely unknown, its allergenicity has been extensively studied. When Fel d 1 is presented by antigen-presenting cells, such as dendritic cells, it triggers the production of IgE antibodies during sensitization. These IgE antibodies specifically target and bind to epitopes on Fel d 1, leading to the activation of immune cells and the subsequent release of inflammatory mediators. This immune response causes the clinical manifestation of allergic symptoms [4]. In addition, T cell epitopes recognized by helper T cells can indirectly induce IgE production through T cell-mediated reactions, further amplifying the allergic response [5]. Thus, the recognition of IgE bound to effector cells, such as basophils and mast cells, plays a crucial role in the allergic cascade [6].

Accurate identification and understanding of epitopes related to allergies are essential for comprehending allergic responses and developing diagnostic tools. Experimental identification of B-cell epitopes has traditionally been the “gold standard” for epitope mapping [7]. However, this approach is often costly, time-consuming, and requires specialized laboratory techniques. Alternatively, computational epitope prediction methods have emerged as practical tools for epitope identification [8]. These methods utilize algorithms and machine learning techniques to analyze the physicochemical properties of proteins and predict potential epitopes.

Early computational tools for B-cell epitope prediction relied on amino acid propensity scales, which characterized the physicochemical properties of B-cell epitopes such as hydropathy [9–10], flexibility [11], and surface accessibility [12]. These scales provided a foundation for initial predictions but had limitations in accuracy and specificity. However, recent advancements in machine learning techniques have leveraged computational B-cell epitope prediction methods, leading to improved outcomes. Machine learning algorithms, including Support Vector Machine (SVM) [13–15], Random Forest (RF) [16–17], K-Nearest Neighbors (KNN) [18], and Manifold Adaptive Experimental Design (MAED) [19], have been developed and trained to differentiate known B-cell epitopes from non-epitopes [14–17, 20–26].

While there have been many computational methods for epitope prediction, their application in predicting IgE and T-cell epitopes, specifically in the context of allergenic proteins has been largely limited. IgE epitopes are of particular interest due to their direct involvement in allergic reactions. However, there have been only a few computational methods developed specifically for IgE epitope prediction [27–28], and most of them are currently unavailable. Therefore, in this study, we aimed to evaluate the effectiveness of currently available general T- and B-cell computational epitope prediction tools and their capacity to accurately identify allergenic epitopes.

To accomplish this, we employed an assortment of epitope prediction tools accessible through the Immune Epitope Database (IEDB) [29] and other sources. Our objective was to perform a detailed assessment of these methods, focusing on their ability to identify Fel d 1 allergenic epitopes.

## Materials and methods

### Epitope prediction tools

To computationally predict Fel d 1 epitopes, we mainly used epitope prediction tools from the Immune Epitope Database (IEDB). The IEDB hosts a comprehensive suite of seven sequence-based B-cell epitope prediction tools (BepiPred-1.0 [17], BepiPred-2.0 [16], Chou and Fashman β-turn prediction [30–31], Emini surface accessibility scale [12], Karplus and Schulz flexibility prediction scale [11], Kolaskar and Tongaonkar antigenicity scale [32], and Parker hydrophilicity scale [33]) along with two structure-based prediction methods (ElliPro [34] and DiscoTope [23]). For the structure-based methods, we utilized the dimeric structure of Fel d 1 (PDB ID: 1PUO). Among the structure-based tools, we specifically chose ElliPro, but excluded DiscoTope, as DiscoTope predicted all residues of Fel d 1 to be epitopes. In addition to the IEDB’s B-cell epitope prediction tools, we also used BepiPred-3.0 [35], a protein language model epitope predictor. We applied default cut-off values for each prediction method.

For the prediction of T-cell epitopes, we employed two pMHC-II binding prediction tools: ProPred [36] and the IEDB MHC-II binding prediction tool [37]. In ProPred, the input sequence is divided into overlapping 9-mer linear peptides. We set a 5% threshold for ProPred to characterize each nonamer as either a binder or a non-binder for 8 HLA types [38]. As for the IEDB prediction method, the query sequence was fragmented into overlapping 15-mer linear peptides, and binding prediction was performed for each peptide using the 27 HLA reference set [39]. In this study, we used the IEDB recommended 2.22 method, using the percentile rank as a binding indicator. Peptides with a rank below 10 were classified as binders, while those with a higher rank were considered non-binders. The epitope scores for each method were calculated as a linear sum of the number of binding events. To evaluate the known T-cell epitopes shorter than 15 residues using the IEDB prediction tool, extra residues at each terminus were added to make the peptides within a 15mer window (**Table 1**).

**Table 1.**
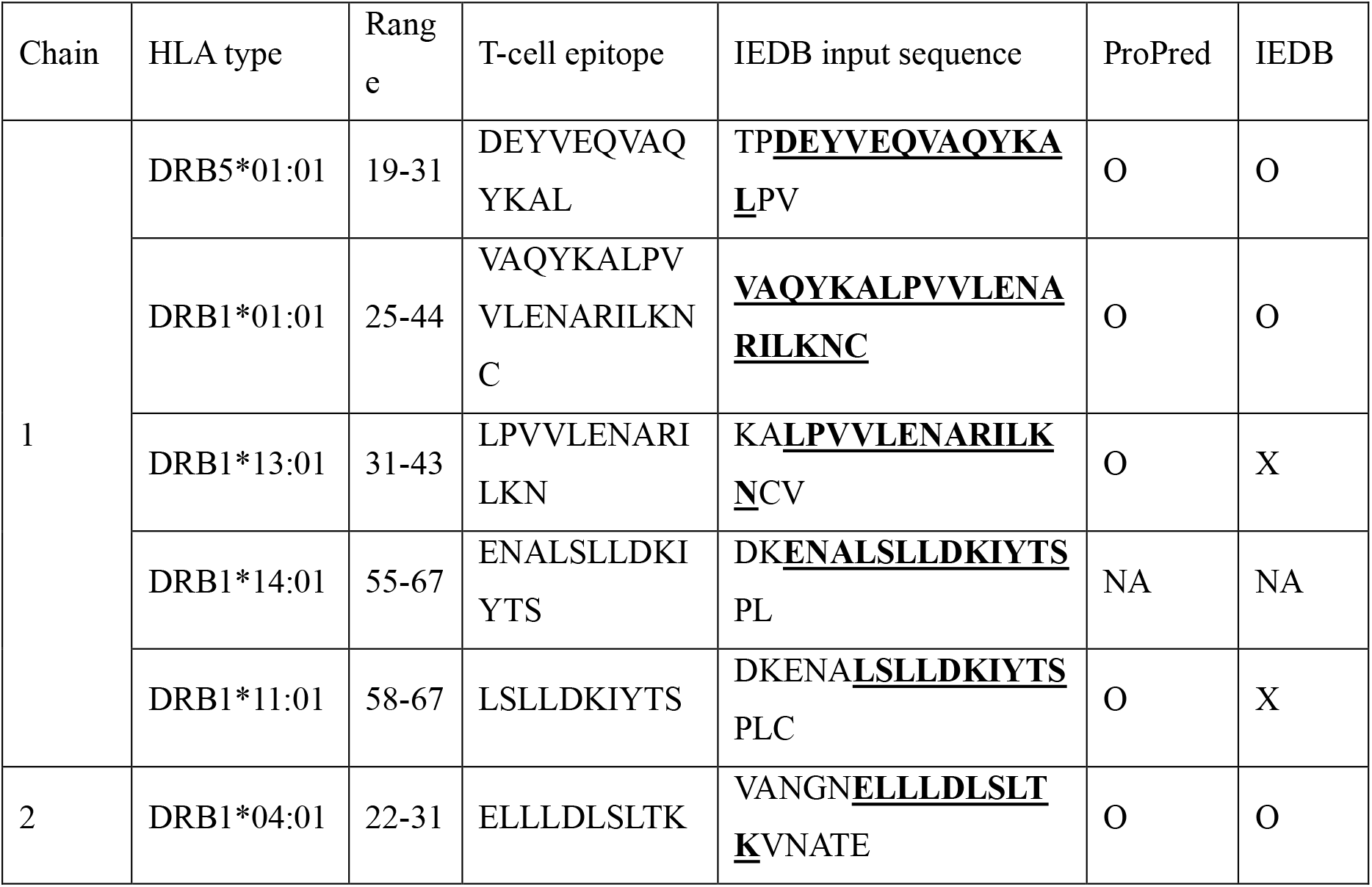
T-cell epitopes of Fel d 1 allergen and their predictions using IEDB and ProPred methods. As the default sequence length for the IEDB T-cell epitope prediction method is 15, we extended the input sequence when its length was shorter than 15 residues. “O” indicates that the method successfully identified the peptide as an epitope for a given allele, while “X” denotes a failure in prediction. Neither of the prediction methods included HLA-DRB1*14:01 in their allele sets and thus no prediction was possible for the specific peptide (ENALSLLDKIYTS).

### Allergen epitope database

To evaluate general predictive abilities of the B-cell epitope prediction methods, we performed benchmarking using the Allerbase database [40]. This database contains a total of 117 allergens and 1,134 linear IgE epitopes. We selected allergens with determined crystal structures from the set, resulting in a subset of 59 allergens with 306 IgE epitopes (**S1 File**). For the T-cell epitope prediction, we used a food allergen database [41], which includes 81 epitopes from 9 allergens (**S1 File**).

### Metrics for epitope/nonepitope binary classification

To evaluate the binary classification of regions as epitope or nonepitope, we employed several statistical metrics: Matthew’s Correlation Coefficient (MCC), Precision (Positive/Negative Predictive values), Recall (Sensitivity/Specificity), and F1 Score.

MCC is a balanced measure that accounts for true and false positives and negatives, ideal for uneven class sizes. It ranges from -1 (total disagreement) to 1 (perfect prediction), with values above 0.5 indicating good predictive performance. Positive Predictive Value (PPV) reflects the accuracy of the model in identifying regions as epitopes, with higher values suggesting better precision. Negative Predictive Value (NPV) indicates the proportion of nonepitope regions correctly identified, with values closer to 1 denoting higher accuracy in predicting nonepitopes. Recall (Sensitivity and Specficity) measures the model’s ability to identify all relevant epitope (nonepitope for specificity) instances correctly, with a range from 0 to 1. The F1 Score is the harmonic mean of Precision (PPV or NPV) and Recall (Sensitivity or Specificity), useful for uneven class distributions, ranging from 0 (worst) to 1 (best).

### Defining Fel d 1 epitopes

While it is known that some Fel d 1 epitopes are partially conformational, our focus in this study was on the linear epitopes of Fel d 1 [42]. This choice was made due to the general low consistency and accuracy of computational prediction methods for conformational epitopes [43].

Fel d 1 is a heterodimer composed of two polypeptide chains (**Fig 1A**), known as Chain 1 and Chain 2 (also referred to as α and β chains, respectively). Chain 1 consists of 70 amino acids and has an approximate molecular mass of 8 kDa. Chain 2 exhibits variation in its C-terminal region, resulting in three known isoforms [2, 44–45]. The major linear IgE-binding epitopes of Fel d 1 are located in three regions: two in Chain 1 and one in Chain 2 (Chain 1: 25 VAQYKALPVVLENA 38, 46 DAKMTEEDKRNALS 59, Chain 2: 12 DVFFAVANGNELLL 25) [46]. T-cell epitopes are present in a total of six regions, including overlapping segments (**Table 1**) [47].

**Fig 1.**
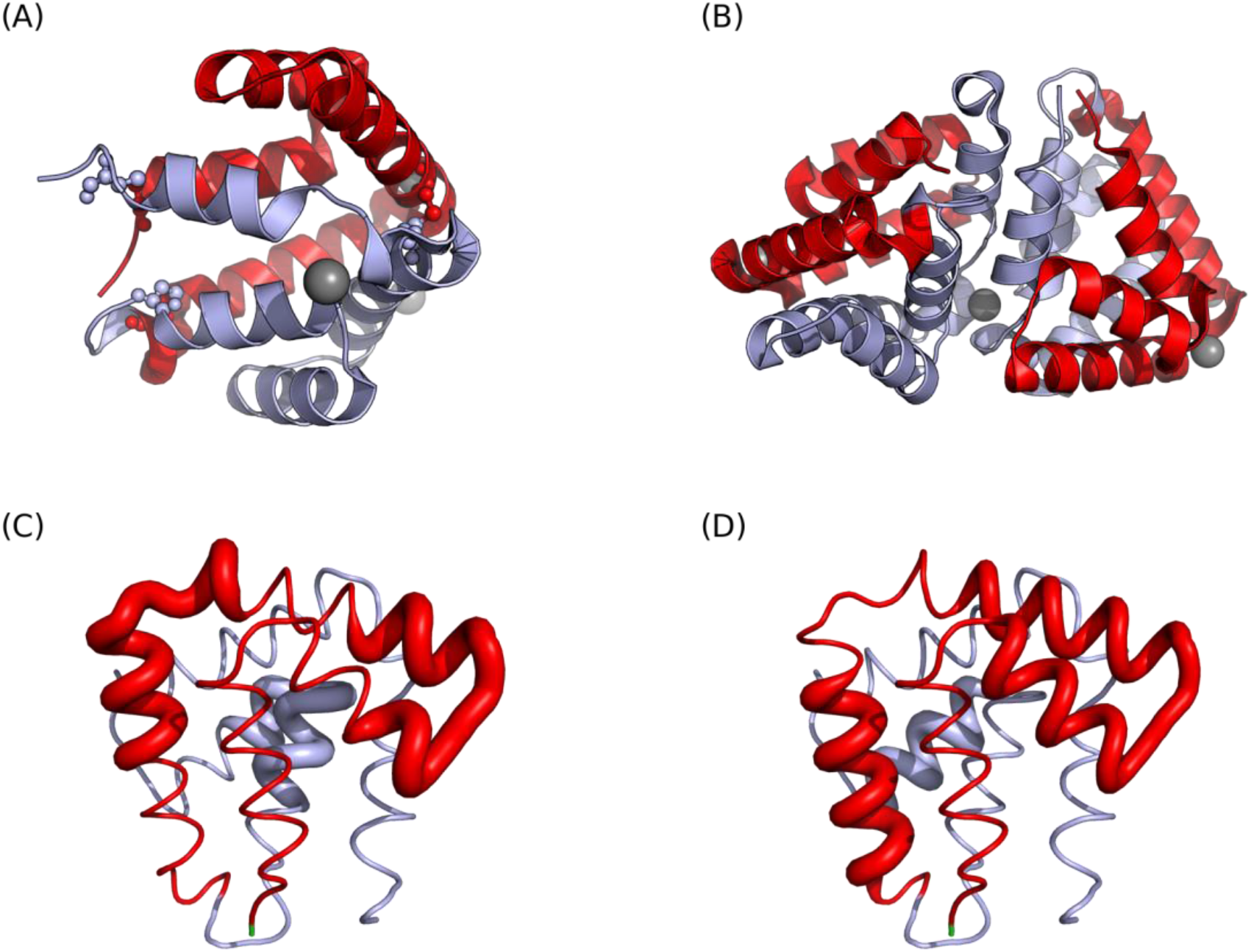
The structures and epitopes of Fel d 1. (A) The heterodimeric structure of Fel d 1 (PDB ID: 1PUO). The three disulfide bonds are highlighted in a small sphere representation. Chain 1 is colored in red and Chain 2 is in skyblue. (B) The tetrameric structure of Fel d 1 (PDB ID: 2EJN). Two Fel d 1 heterodimers are non-covalently bonded between two Chain 2s. (C) Three IgE epitopes of Fel d 1. (D) The residues that are included in the six T-cell epitopes (see **Table 1**). Overall, there are three main T-cell epitope regions. Epitopes are represented in thicker ribbons in (C) and (D).

## Results

### Structure analysis of Fel d 1 epitopes

Fel d 1 primarily adopts the alpha-helical heterodimeric structure (**Fig 1A**). The heterodimer is composed of two distinct chains (Chain 1 and 2) that are linked by three disulfide bonds. These heterodimers naturally assemble into tetramers via non-covalent interactions between Chain 2s, as shown in **Fig 1B** [48–50]

Given the structure and formation of the tetramer, the IgE epitope located on Chain 2 might be less accessible for IgE binding when the protein is in its tetrameric structure (The solvent accessible surface area for the epitope is 1390.99 Å^2^ in dimer and 381.83 Å^2^ in tetramer). It is known that approximately 15 % of Fel d 1 is tetrameric in house dust [51]. As the binding of IgE antibodies to their specific epitopes is a crucial step in the onset of allergic reactions, the non-covalent interactions between Chain 2s in the tetramer might render these epitopes less exposed, thereby decreasing the likelihood of IgE binding.

### Prediction of IgE epitopes

To assess the IgE epitope prediction, we examined nine publicly available epitope prediction tools, which include web-based B-cell epitope prediction methods provided by the IEDB. Generally, most methods are better at predicting non-epitopes than epitopes for Fel d 1 (**Fig 2 and Table 2**). However, considering the imbalance between the number of epitopes and non-epitopes, with 42 out of 162 total residues being epitopes (28 out of 70 in Chain 1, and 14 out of 92 in Chain 2), the probability of correctly predicting epitopes by chance alone is significantly lower (0.26) compared to non-epitopes (0.74). The Matthews correlation coefficient (MCC) values suggest that most methods perform similarly to random predictions.

**Fig 2.**
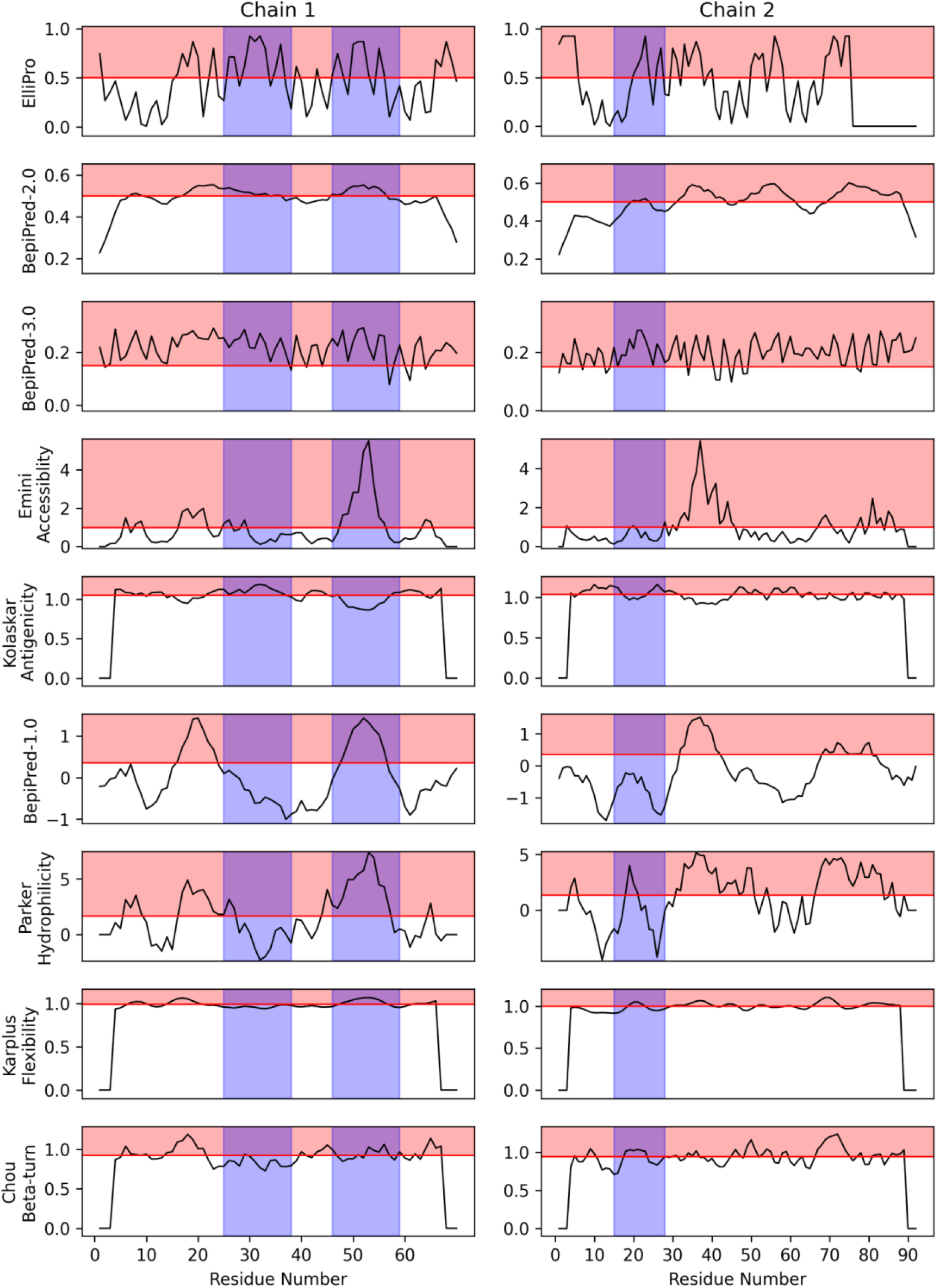
IgE epitope prediction using computational tools. Horizontal red lines represent the threshold values used by each prediction method. Red shaded areas indicate regions predicted as epitopes. Blue shaded areas denote the actual epitope positions of Fel d 1. Most methods are essentially no different from random selection. See also **Table 2** for other details.

**Table 2.** Prediction results of computational B-cell epitope prediction tools for Fel d 1 IgE epitopes. Overall, the majority of methods yield results that are indistinguishable from random selection. See also **Fig 2** for other details.

|  | MCC | Prediction of Epitopes |  |  | Prediction of Non-epitopes |  |  |
| --- | --- | --- | --- | --- | --- | --- | --- |
|  |  | PPV | Sensitivity | F1 | NPV | Specificity | F1 |
| **ElliPro** | 0.218 | 0.34 | 0.48 | 0.4 | 0.79 | 0.68 | 0.73 |
| **BepiPred-2.0** | 0.147 | 0.32 | 0.64 | 0.43 | 0.81 | 0.53 | 0.64 |
| **BepiPred-3.0** | 0.136 | 0.28 | 0.95 | 0.43 | 0.9 | 0.15 | 0.26 |
| **Emini** | 0.016 | 0.27 | 0.33 | 0.3 | 0.75 | 0.68 | 0.71 |
| **Kolaskar** | 0.034 | 0.27 | 0.55 | 0.37 | 0.76 | 0.49 | 0.6 |
| **BepiPred-1.0** | -0.053 | 0.22 | 0.21 | 0.22 | 0.73 | 0.73 | 0.73 |
| **Parker** | -0.013 | 0.25 | 0.45 | 0.32 | 0.74 | 0.53 | 0.62 |
| **Karplus** | -0.125 | 0.19 | 0.31 | 0.24 | 0.69 | 0.55 | 0.61 |
| **Chou** | -0.090 | 0.22 | 0.38 | 0.28 | 0.7 | 0.52 | 0.6 |

BepiPred-3.0 excels at identifying true non-epitopes (NPV 0.9) but also produces considerable false positives in epitope predictions and misses many true non-epitopes (Specificity 0.15). BepiPred-2.0 and ElliPro show a more balanced performance between precision and recall for Fel d 1, with ElliPro having a slight edge in F1 score.

In an effort to extend our analysis beyond Fel d 1, we assessed the prediction accuracy of the same nine epitope prediction tools on 59 different allergenic proteins. Out of a total of 12,046 residues, 3,521 were allergen epitopes, comprising approximately 29 % of the total epitopes. Consistent with the results observed in the case of Fel d 1, the prediction tools generally exhibited better performance in predicting non-epitopes compared to epitopes (**S1 Table**). However, most tools exhibit MCC values close to 0, indicating no significant difference from random predictions. Our results indicate that the effectiveness of the current B-cell epitope prediction tools is largely limited, highlighting the need for continuous refinement.

### Prediction of T-cell epitopes

T-cell immunity plays a pivotal role in managing allergic responses by dictating the magnitude and nature of the immune response to allergens. In particular, T-cell responses against allergens can have profound influence on the production of specific types of antibodies, such as IgE, which plays a direct role in allergic reactions.

In our initial investigations, we subjected the known Fel d 1 T-cell epitopes to ProPred and the IEDB prediction methods. Among these epitopes, the sequence 55 ENALSLLDKIYTS 67 in Chain 1, known for its binding to HLA-DRB1*14:01, was unable to be evaluated since the specific allele is not included in either of the methods. Notably, we found that the IEDB method was not able to correctly predict two of the known T-cell epitopes as binders for their respective alleles. Contrarily, ProPred correctly identified all the epitopes as binders for their associated alleles (**Table 1**).

To explore the general predictive abilities of ProPred and IEDB for T-cell epitopes across various allergens, we conducted the same analyses on a total of 81 T-cell epitopes from 9 allergens (**S2 Table**). Our benchmark reveals that the two predictors exhibit different tendencies in prediction at the cut-off values we used. While ProPred is better at predicting non-epitopes compared to the IEDB method (F1 score: 0.84 vs. 0.65), IEDB is slightly better than ProPred for epitope prediction (F1: 0.18 vs. 0.31). However, the MCC values show that both methods are only marginally better than random (MCC: 0.04 vs. 0.11).

These observations are consistent when predicting T-cell epitopes on Fel d 1. Applying both methods to full sequences of Chain 1 and 2, ProPred remains superior at predicting non-epitopes (F1: 0.87 vs. 0.6), although the MCC values for both are only weakly positive (0.27 vs. 0.15, see **Fig 3 and Table 3**).

**Fig 3.**
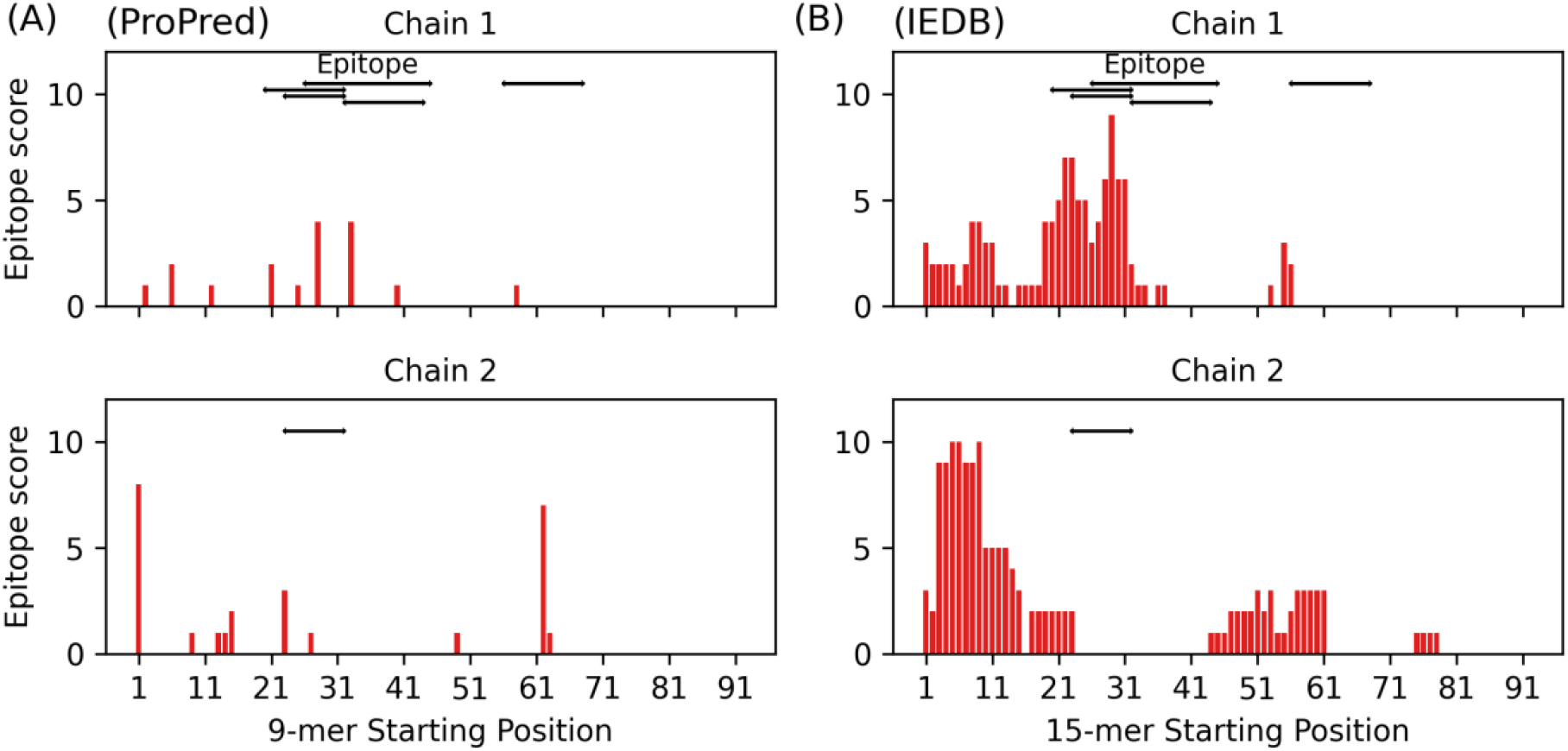
Fel d 1 T-cell epitope maps and prediction results. Black horizontal lines represent known T-cell epitope positions, while vertical red lines denote the predicted epitope scores of each peptide starting at these positions. Higher scores indicate a greater number of binding events. In general, ProPred is better at predicting non-epitopes, while both methods perform only marginally better than random selection. Additional details are provided in **Table 3** and **S1 File**.

**Table 3.** Prediction results of computational pMHC-II binding prediction tools for Fel d 1 epitopes. ProPred demonstrates enhanced accuracy in predicting non-epitopes. However, according to the Matthews Correlation Coefficient (MCC) values, the overall prediction results are only slightly better than random. See also **Fig 3** and **S1 File** for other details.

|  | MCC | Prediction of Epitopes |  |  | Prediction of Non-epitopes |  |  |
| --- | --- | --- | --- | --- | --- | --- | --- |
|  |  | PPV | Sensitivity | F1 | NPV | Specificity | F1 |
| **IEDB** | 0.272 | 0.17 | 1 | 0.29 | 1 | 0.43 | 0.6 |
| **ProPred** | 0.148 | 0.32 | 0.29 | 0.27 | 0.85 | 0.89 | 0.87 |

It is important to note that these methods solely predict peptide-MHC binding events. High epitope binding scores do not necessarily indicate immunogenicity, as actual immune responses also require T-cell recognition. Consequently, the absence of pMHC binding typically means no immune response, while its presence does not guarantee an immune response. Therefore, although the prediction methods are nearly random or only marginally better than random, the high F1 score for non-epitope prediction by ProPred may be beneficial for reducing T-cell epitope content [52–53].

## Discussion

In our study, we assessed various computational methods for predicting Fel d 1 epitopes, demonstrating their usefulness and limitations in allergenic epitope identification. Though focused on Fel d 1, these tools can be utilized for other allergens, potentially informing allergen-specific therapy and diagnostics development.

While these computational methods show promise in other applications, they fall short in accurately identifying allergenic epitopes, typically predicting non-epitopes more effectively. Most B-cell epitope prediction methods are essentially no different from random selection, possibly due to their lack of specific design for IgE epitopes.

For T-cell epitope prediction, ProPred successfully identified all known T-cell epitopes for the associated alleles in our analysis. In contrast, the IEDB method failed to predict two known T-cell epitopes. While ProPred is generally better at predicting non-epitopes, overall predictive abilities of the two methods are only marginally better than random.

In conclusion, our study highlights the limitations of current computational methods in allergenic epitope prediction. Particularly for B-cell epitope prediction, the modest improvement in MCC values between the two BepiPred versions suggests that integrating advanced artificial intelligence techniques and 3D structure information could enhance the accuracy and specificity in the future. Expansive and comprehensive databases of experimentally validated allergen epitopes could therefore assist these tools in refining their algorithms, thereby improving the prediction of epitopes.

## Acknowledgments

This study was financially supported by Chonnam National University (Grant number: 2021-2216) and National Research Foundation of Korea grants funded by the Korean government (NRF-2020R1A5A2031185, NRF-2020M3A9G3080281, and 2021R1F1A1063769).

## Supporting information

**S1 Table.** Prediction results of a computational B cell epitope prediction tool for known allergen IgE epitopes.

|  | MCC | Prediction of Epitopes |  |  | Prediction of Non-epitopes |  |  |
| --- | --- | --- | --- | --- | --- | --- | --- |
|  |  | PPV | Sensitivity | F1 | NPV | Specificity | F1 |
| **ElliPro** | 0.035 | 0.29 | 0.52 | 0.38 | 0.74 | 0.52 | 0.61 |
| **BepiPred-2.0** | 0.058 | 0.32 | 0.52 | 0.39 | 0.74 | 0.55 | 0.63 |
| **BepiPred-3.0** | 0.129 | 0.36 | 0.46 | 0.41 | 0.76 | 0.67 | 0.71 |
| **Emini** | 0.049 | 0.32 | 0.39 | 0.35 | 0.73 | 0.66 | 0.7 |
| **Kolaskar** | 0.003 | 0.29 | 0.33 | 0.31 | 0.71 | 0.67 | 0.69 |
| **BepiPred-1.0** | -0.010 | 0.28 | 0.41 | 0.33 | 0.71 | 0.58 | 0.64 |
| **Parker** | -0.032 | 0.29 | 0.61 | 0.4 | 0.72 | 0.41 | 0.52 |
| **Karplus** | 0.014 | 0.28 | 0.52 | 0.36 | 0.69 | 0.45 | 0.54 |
| **Chou** | 0.015 | 0.29 | 0.67 | 0.41 | 0.72 | 0.34 | 0.46 |
| **Random** | 0.097 | 0.3 | 0.32 | 0.29 | 0.68 | 0.77 | 0.71 |
| **Number of residues** |  | 3,553 (29 %) |  |  | 8,807 (71%) |  |  |

**S2 Table.**
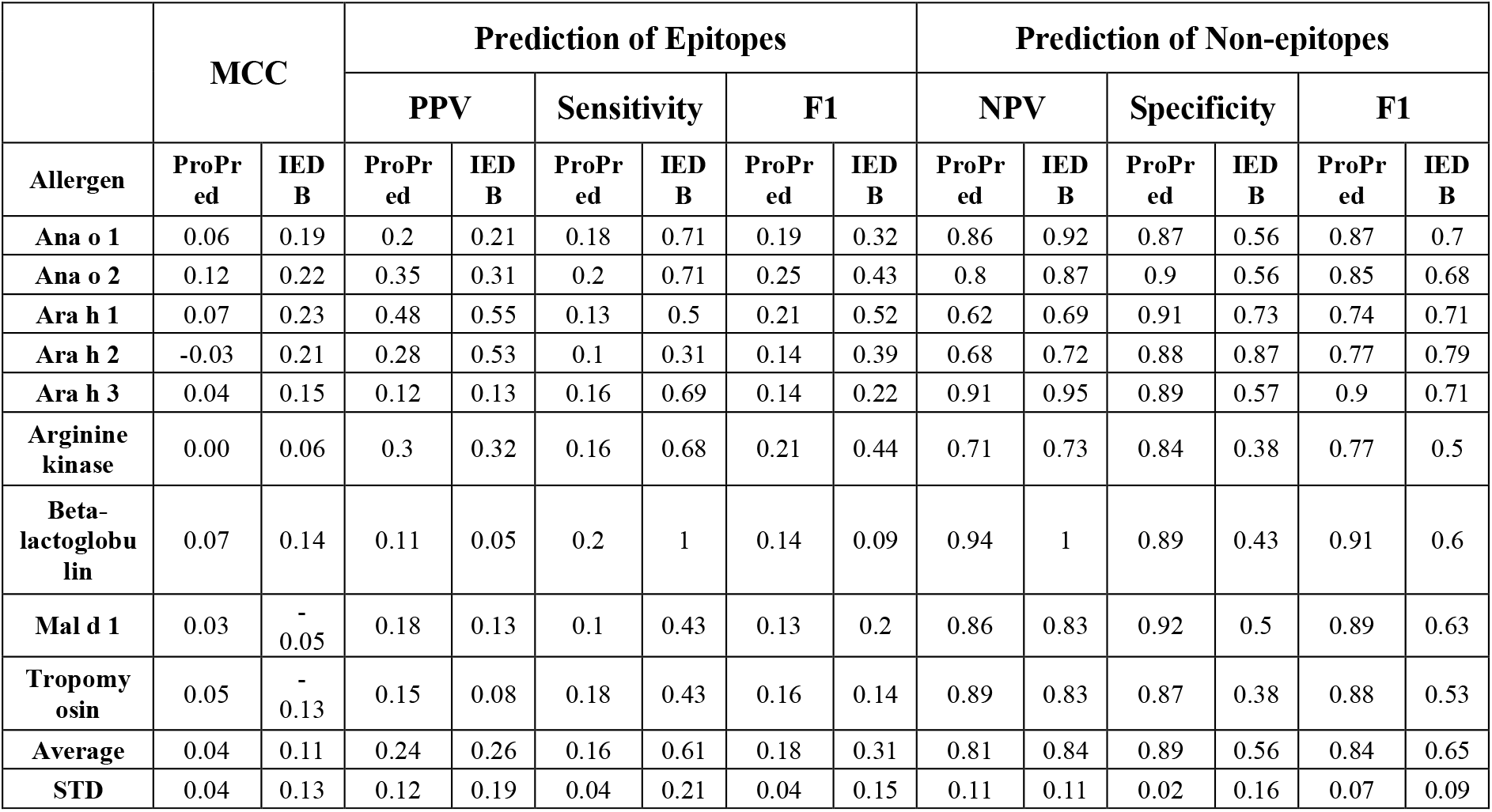
T-cell epitope prediction results for food allergen IgE epitopes using ProPred and IEDB prediction methods.

|  | MCC |  | Prediction of Epitopes |  |  |  |  |  | Prediction of Non-epitopes |  |  |  |  |  |
| --- | --- | --- | --- | --- | --- | --- | --- | --- | --- | --- | --- | --- | --- | --- |
|  |  |  | PPV |  | Sensitivity |  | F1 |  | NPV |  | Specificity |  | F1 |  |
| Allergen | ProPred | IEDB | ProPred | IEDB | ProPred | IEDB | ProPred | IEDB | ProPred | IEDB | ProPred | IEDB | ProPred | IEDB |
| Ana o 1 | 0.06 | 0.19 | 0.2 | 0.21 | 0.18 | 0.71 | 0.19 | 0.32 | 0.86 | 0.92 | 0.87 | 0.56 | 0.87 | 0.7 |
| Ana o 2 | 0.12 | 0.22 | 0.35 | 0.31 | 0.2 | 0.71 | 0.25 | 0.43 | 0.8 | 0.87 | 0.9 | 0.56 | 0.85 | 0.68 |
| Ara h 1 | 0.07 | 0.23 | 0.48 | 0.55 | 0.13 | 0.5 | 0.21 | 0.52 | 0.62 | 0.69 | 0.91 | 0.73 | 0.74 | 0.71 |
| Ara h 2 | -0.03 | 0.21 | 0.28 | 0.53 | 0.1 | 0.31 | 0.14 | 0.39 | 0.68 | 0.72 | 0.88 | 0.87 | 0.77 | 0.79 |
| Ara h 3 | 0.04 | 0.15 | 0.12 | 0.13 | 0.16 | 0.69 | 0.14 | 0.22 | 0.91 | 0.95 | 0.89 | 0.57 | 0.9 | 0.71 |
| Arginine kinase | 0.00 | 0.06 | 0.3 | 0.32 | 0.16 | 0.68 | 0.21 | 0.44 | 0.71 | 0.73 | 0.84 | 0.38 | 0.77 | 0.5 |
| Beta-lactoglobulin | 0.07 | 0.14 | 0.11 | 0.05 | 0.2 | 1 | 0.14 | 0.09 | 0.94 | 1 | 0.89 | 0.43 | 0.91 | 0.6 |
| Mal d 1 | 0.03 | -0.05 | 0.18 | 0.13 | 0.1 | 0.43 | 0.13 | 0.2 | 0.86 | 0.83 | 0.92 | 0.5 | 0.89 | 0.63 |
| Tropomyosin | 0.05 | -0.13 | 0.15 | 0.08 | 0.18 | 0.43 | 0.16 | 0.14 | 0.89 | 0.83 | 0.87 | 0.38 | 0.88 | 0.53 |
| Average | 0.04 | 0.11 | 0.24 | 0.26 | 0.16 | 0.61 | 0.18 | 0.31 | 0.81 | 0.84 | 0.89 | 0.56 | 0.84 | 0.65 |
| STD | 0.04 | 0.13 | 0.12 | 0.19 | 0.04 | 0.21 | 0.04 | 0.15 | 0.11 | 0.11 | 0.02 | 0.16 | 0.07 | 0.09 |

**S1 File. Comprehensive Dataset of IgE and T-cell Epitope Predictions for Allergen Research.**

## Notes

### Competing Interest Statement

The authors have declared no competing interest.

### Summary of Updates

The benchmark results have been thoroughly re-evaluated.

